# General purpose genotypes and evolution of higher plasticity in clonality underlie knotweed invasion

**DOI:** 10.1101/2024.10.13.618084

**Authors:** Shengyu Wang, Zhiyong Liao, Peipei Cao, Marc W. Schmid, Lei Zhang, Jingwen Bi, Stacy B. Endriss, Yujie Zhao, Madalin Parepa, Wenyi Hu, Hikaru Akamine, Jihua Wu, Ruiting Ju, Oliver Bossdorf, Christina L. Richards, Bo Li

**Author notes:** Correspondence Christina L. Richards and Bo Li should be considered joint senior author., Christina L. Richards, Bo Li, (86) 0871-65933840. Shengyu Wang and Zhiyong Liao should be considered joint first author.

## Abstract

Many widespread invasive plant species express high phenotypic variation across novel environments, providing a unique opportunity to examine ecological and evolutionary dynamics under global change. However, biogeographical studies often lack information about the origin of introduced populations, limiting our understanding of post-introduction evolution of introduced species in the new range. Here, we assessed the responses of *Reynoutria japonica* from 128 populations spanning three latitudinal transects in the native ranges of China and Japan, and the introduced ranges of North America and Europe. When grown in two common gardens in the native range, plants from introduced populations in North America and Europe differed in almost all traits from those from native Chinese populations, but were similar to plants from native Japanese populations. Compared to Chinese populations, North American, European and Japanese populations expressed lower trait values and plasticity in most traits. However, plants from both introduced ranges and from Japanese populations expressed higher clonality and plasticity in clonality than plants from Chinese populations. In addition, introduced populations expressed higher plasticity in clonality but lower plasticity in basal diameter compared to Japanese populations. Our study showed heritable differences in phenotypes between plants from the native ranges of Japan and China, and those from the introduced ranges were similar to the putative source of origin in Japan. However, we found that introduced populations may have evolved higher plasticity in clonal growth. Our findings emphasize the critical role of clonality and plasticity in invasion success, demonstrating the importance of discriminating between source and non-source native populations to identify ecological and evolutionary responses of invasive plants to novel environments.

## 1 INTRODUCTION

Non-native species often experience a novel combination of environmental and reproductive barriers in their introduced ranges (Theoharides & Dukes, 2007), providing a unique opportunity to examine rapid ecological and evolutionary dynamics under global change. Understanding the mechanisms that underlie invasion success is also a global imperative, considering that invasive species are “main direct drivers of biodiversity loss across the globe” (Convention on Biological Diversity, 2022). Many hypotheses have proposed that environmental changes in the introduced range can influence trait expression, potentially exposing novel trait combinations and driving rapid adaptive evolution (Gioria et al. 2023).

Phenotypic plasticity is thought to play an important role in this context either through selection for “general purpose genotypes” (Baker, 1965; Bossdorf et al., 2005; Richards et al., 2006) or as a mechanism to expose cryptic genetic variation that can be refined through genetic accommodation (i.e., the so-called “plasticity first” hypothesis; Levis & Pfennig, 2016). Many studies have shown that introduced populations have greater plasticity than their native conspecifics (Bhattarai et al., 2017; Yang et al., 2021), but there is a lack of clear support that such plasticity can be translated into fitness benefits (Castillo et al., 2021; Boyd et al., 2022). As in other studies of plant invasion, an important limitation to support the roles of plasticity in introduced populations is the identification of an appropriate comparison (van Kleunen et al., 2010; Levis & Pfennig, 2016). Comparing introduced populations with their source native populations could improve our understanding of the roles of plasticity in the success of species invasions (Valladares et al., 2014; Levis & Pfennig, 2016; Hierro et al., 2022).

Studying post-introduction evolution is further complicated by the fact that some species have large geographical distributions in both native and introduced ranges, requiring careful consideration of the invasion history and choice of locations for common garden experiments (Colautti and Lau, 2015; van Boheemen et al., 2019). A growing number of biogeographic studies of native and introduced populations have shown that similar patterns of abiotic stresses shape parallel clines in plant traits in both native and introduced populations (Maron et al., 2004; van Boheemen et al., 2019; Chen et al., 2023). Some studies, on the contrary, have found no similarity between clines in native and introduced populations of a single species (Alexander et al., 2012; Endriss et al., 2018; Liu et al., 2020). There are still many open questions about how adaptation of introduced plants differs between continents and during range expansion, and addressing these questions requires a biogeographic approach (Gioria et al., 2023).

To explore differences in plasticity between native and introduced populations, many studies have manipulated one or a few abiotic factors such as light (Flory et al., 2011), nutrient (Luo et al., 2019), water availability (Liao et al., 2020) or temperature (Molina-Montenegro & Naya, 2012) in a single common garden. However, the conclusions from such studies depend, at least partly, on where the common garden was located as different sites represent different combinations of diverse biotic and abiotic factors (Maron et al., 2007; Woods et al., 2012; Yang et al., 2021). In addition, asexual clonal growth is envisaged as an important trait that facilitates plant invasions (Atwood & Meyerson, 2011; Wang et al., 2017), but evidence of increased clonality has rarely been examined (Bock et al., 2018). Consideration of both plasticity and clonality across broad biogeographical ranges may therefore help improve our understanding of several hypotheses (Felker-Quinn et al., 2013; Liu et al., 2020).

In previous work, we found that plants in introduced populations of *Reynoutria japonica* grew larger and had higher nutritional value compared to conspecific plants in native populations (Irimia et al., 2024). Several other studies have reported that introduced populations of *R*. *japonica* consisted of a single genotype in Europe (Hollingsworth & Bailey, 2000; Zhang et al., 2016) and the USA (Richards et al., 2012; Gaskin et al., 2014; Groeneveld et al., 2014). Here, we aimed to clarify the roles of phenotypic plasticity and rapid evolution within the context of the introduction of such a putative “general purpose genotype”. A “general purpose genotype” should be characterized by high phenotypic plasticity, particularly in traits that contribute to invasion success (Baker, 1965; Richards et al., 2006). Spread of such genotypes could also be enhanced by clonality (Baker, 1965), and initial plasticity in clonality could expose variation among individuals in the ability to respond with clonal growth (i.e., “plasticity first”). In that case, we might expect selection for increased plasticity in clonality (genetic accommodation of plasticity, Bock et al., 2018).

To evaluate the contributions of plasticity and clonality to invasion success, we compared growth of *R. japonica* from 55 native-range populations (Japan and China) and 73 introduced-range populations (North America and Europe) in two common gardens in China. We tested several alternative hypotheses: (1) plants from introduced ranges exhibit increased growth compared to those from native ranges when grown in common gardens; (2) responses of the introduced populations are more similar to the native Japanese populations (the putative source of the introduced plants) than to the native Chinese populations; (3) the introduced populations have evolved greater phenotypic plasticity than the native populations.

## 2 MATERIALS AND METHODS

### 2.1 Study species and collecting sites

*Reynoutria japonica* (Houtt.) Ronse Decraene (Japanese knotweed, Polygonaceae) is native to eastern Asia (Bailey & Conolly, 2000) and was introduced into Europe in the mid-19^th^ century (Beerling et al., 1994; Bailey & Conolly, 2000) and into North America before 1873 (Barney, 2006). Because of the highly negative effects of *R. japonica*, it has been listed as one of the 100 world’s worst invasive species by the IUCN (Lowe et al., 2000). Introduced populations of *R. japonica* have almost no genetic diversity, and are thought to originate from a single female individual (Hollingsworth & Bailey, 2000; Richards et al., 2012; Gaskin et al., 2014; Groeneveld et al., 2014; Jugieau et al., 2024).

From 2019 to 2020, we conducted a cross-latitudinal survey of 150 sites in the native range of China and the introduced ranges of North America and Europe (Figure 1, Irimia et al., 2024). Briefly, we collected rhizomes from 5 individuals at each of 50 populations along a 2000 km transect in each range, with each population approximately 40 km apart from neighboring populations (see more details in Irimia et al., 2024). In addition, we collected rhizomes from 6 populations in Nagasaki, Japan separated by about 20 km because this area was known as the potential source of North American and European populations (Beerling et al., 1994; Bailey & Conolly, 2000). We collected 5–8 individuals from each Japanese population between 28 April and 4 May 2021. Our chloroplast sequencing data also showed that the Japanese populations we collected share the same chloroplast haplotype as the introduced North American and European populations (Zhang et al., under review). We therefore considered the Japanese populations as the source within the native range.

**Figure 1.**
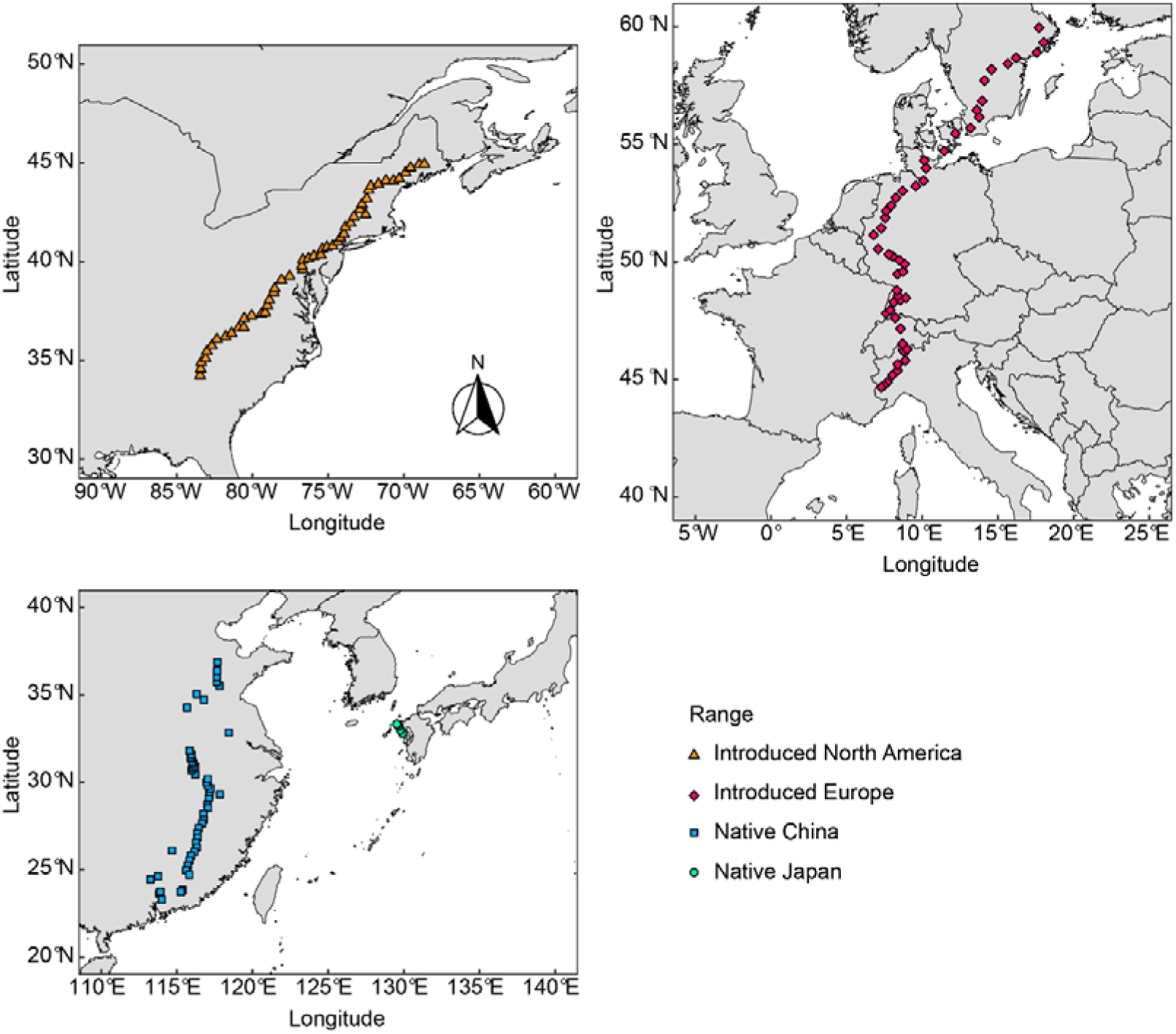
Locations of *R. japonica* populations sampled in the field. 50 populations along latitudinal transects were sampled in the introduced North American range, introduced European range, and native Chinese range. 6 populations were sampled in native Japanese range from the area that is considered to be the source of the introduced populations.

We transplanted rhizome fragments to a quarantine greenhouse in Xishuangbanna Tropical Botanical Garden, Chinese Academy of Sciences (Mengla, Yunnan, China). Due to import limitations, we obtained only one individual from each North American population. Because of variable cultivation success during the experimental period, we ended up with a total of 464 *R. japonica* individuals from 55 native populations (50 from China, 5 from Japan) and 73 introduced populations (27 from North America, 46 from Europe) (Table S1). For most of these individuals, we cut two pieces of rhizomes each with at least one bud to ensure their survival and rhizome size did not have significant size effects on plant traits (Figure S1). We removed the fine roots to minimize maternal effects and stored the rhizomes at 4 °C for at least 4 weeks to facilitate sprouting success.

### 2.2 Experimental set-up

We established two common gardens: one at Xishuangbanna Tropical Botanical Garden, Chinese Academy of Sciences, Yunnan (XTBG; 101°16′ E, 21°54′ N) and the other at Fudan University, Shanghai (SH; 121°30′ E, 31°20′ N). The XTBG garden has a tropical monsoon climate, and is close to the southern limit of the native range. The SH garden has a moderate temperate-zone climate, and is roughly at the distributional center of the native range.

We used identical plant cultivation methods in each garden (also see Cao et al., 2024), including potting substrates and fertilization. On 11 March 2022, we treated all the rhizomes with fungicide and transplanted them separately into outdoor 20 L plastic pots with standard soil (Pindstrup substrate 5–20 mm, Pindstrup Mosebrug A/S, Denmark). We randomly arranged all pots into five blocks within each garden, with one individual from each population in each block for European and Chinese populations. For the Japanese and the North American populations, we randomly assigned an even number of pots into each of the blocks (5-7 in each block). The distance between neighbor pots was at least 90 cm to avoid aboveground interference. We added 10 g Osmocote fertilizer (Osmocote plus 801, N: P: K 16:8:12, Everris International B.V., Heerlen, Netherlands) onto the soil surface in each pot at the beginning and mid-point of the experiment. Throughout the experiment, we watered the plants when needed. We put a saucer under each pot to avoid water and nutrient loss. Due to limited rhizome availability and growth success for some individuals, we eventually measured trait values of 464 individuals of *R. japonica* at the XTBG garden (233 individuals from 50 Chinese populations, 25 individuals from 5 Japanese populations, 190 individuals from 46 European populations, and 16 individuals from 16 North American populations) and 519 individuals at the SH garden (240 from 50 Chinese populations, 34 from 5 Japanese populations, 218 from 46 European populations, and 27 from 27 North American populations).

### 2.3 Measurements

In July 2022, we counted the number of ramets (our estimate of clonality) that were > 5 cm tall in each pot. We also scanned the top 5 fully expanded leaves of the tallest ramet (Epson Perfection V550, Japan), and calculated the total leaf area of the 5 leaves after subtracting any area missing due to herbivory damage (with Adobe Photoshop using ImageJ, Schneider et al., 2012) as leaf size. In October 2022, we measured the basal diameter and height of the tallest ramet in each pot, as most of the plants had turned yellow and had reached the end of growing season. We then harvested all aboveground parts and dried all plant materials at 75 °C for at least 72 h before determination of dry mass. And the data are available online (Wang et al., 2024).

### 2.5 Statistical analyses

We performed all analyses in R 4.3.2 (R Core Team 2023). To visualize the overall differences in plant trait values among ranges, we used two separate Horn’s parallel analyses to perform principal component analyses (PCA) on all measured traits in each of the two common gardens (Revelle, 2024). For each PCA, we used the mean trait value within a garden for each population (i.e., one data point per population). To evaluate differences in trait means and plasticities among ranges, we used linear models and all data from the four ranges in the two gardens (i.e., 983 plants). We fitted factors sequentially (type I sum of squares) following general design principles (Schmid et al., 2017) as indicated in Table 1. Significance tests were based on F tests using appropriate error terms and denominator degrees of freedom (Schmid et al., 2017). The fixed terms of the models were garden, range, and their interactions, the random terms were population and the interaction between garden and population. The factor “range” was further split into three contrasts with one degree of freedom each (Schmid et al., 2017): The first compared CN to the other three ranges, the second compared JA to EU and US, and the third compared EU to US. Given that population was treated as a random term, the factor range and its contrasts were compared to the factor population. Likewise, the interaction of the factor range and its contrasts with the factor garden were compared to the interaction between population and garden. In this model, the main effect of the factor range and its contrast compared trait means, and the interactions compared trait plasticities. To compare all pairwise-differences in trait means, we used the same model structure to run linear mixed models implemented in the *lme4* package (Bates et al., 2015), along with Tukey’s Honestly Significant Difference tests (Tukey’s HSD) implemented in the *emmeans* package (Lenth, 2022).

**Table 1.** The effects of range, garden and their interactions on plant traits of *R. japonica* shown as the percent sum of explained by each term. Statistical significance is indicated as: ns, *P* ≥ 0.1; #, *P* < 0.1; *, *P* < 0.05; **, *P* < 0.01; ***, *P* < 0.001. Above. mass, aboveground biomass; Df, Degrees of freedom. *P*-values were calculated by treating population as a random factor. Thus, the factor range and its contrasts were tested against population, and the interactions of range and its contrasts with garden against garden:population. Garden was tested against the residuals. Even though population was treated as a random factor, we provided the variation explained by it and the results of a test against residuals.

| Term | Df | Height | Diameter | Leaf size | Leaf mass | Stem mass | Above. mass | No. ramets |
| --- | --- | --- | --- | --- | --- | --- | --- | --- |
| Garden | 1 | 26.73 *** | 53.69 *** | 4.18 *** | 78 *** | 64.31 *** | 81.17 *** | 5.97 *** |
| Range, contrasts: | 3 | 39.49 *** | 15.85 *** | 31.15 *** | 0.51 # | 9.18 *** | 0.24 ns | 51.33 *** |
| a) CN vs. JA/EU/US | 1 | 39.38 *** | 15.4 *** | 30.72 *** | 0.5 * | 9.16 *** | 0.24 # | 51.13 *** |
| b) JA vs. EU/US | 1 | 0.02 ns | 0.4 * | 0 ns | 0 ns | 0.01 ns | 0 ns | 0.13 # |
| c) EU vs. US | 1 | 0.08 ns | 0.05 ns | 0.43 ns | 0 ns | 0.01 ns | 0 ns | 0.07 ns |
| Population | 123-124 | 9.43 *** | 7.89 | 21.59 *** | 9.01 *** | 11.26 *** | 8.85 *** | 5.07 ns |
| Garden:Range, contrasts: | 3 | 0.28 ns | 3.85 *** | 2.13 *** | 0.08 ns | 3.97 *** | 0.34 ** | 7.34 *** |
| a) Garden:CN vs. JA/EU/US | 1 | 0.19 * | 3.55 *** | 1.75 *** | 0.03 ns | 3.97 *** | 0.31 *** | 6.65 *** |
| b) Garden:JA vs. EU/US | 1 | 0.09 ns | 0.29 *** | 0.02 ns | 0.03 ns | 0 ns | 0.02 ns | 0.68 *** |
| c) Garden:EU vs. US | 1 | 0 ns | 0.01 ns | 0.36 # | 0.02 ns | 0 ns | 0 ns | 0 ns |
| Garden:Population | 112-113 | 4.89 *** | 2.93 # | 10.81 *** | 2.92 *** | 3.13 *** | 2.47 *** | 2.11 ns |
| Residuals | 707-738 | 19.19 | 15.78 | 30.14 | 9.49 | 8.16 | 6.92 | 28.18 |

## 3 RESULTS

We found that on average the trait means and plasticity of introduced populations differed from those of native Chinese populations but were very similar to Japanese populations. An important exception was the increased plasticity in clonality in introduced ranges compared to Japanese populations.

### 3.1 Plant trait divergence among ranges

The first two components of the PCA explained 80.1% and 85.3% of the trait variation across all four ranges for plants grown in the Xishuangbanna (Figure 2a) and Shanghai (Figure 2b) common gardens, respectively. The PCA clearly showed that plants from populations in China separated from those in the other three ranges in both gardens. Plants from Japan, North America and Europe largely overlapped in both gardens in multivariate space.

**Figure 2.**
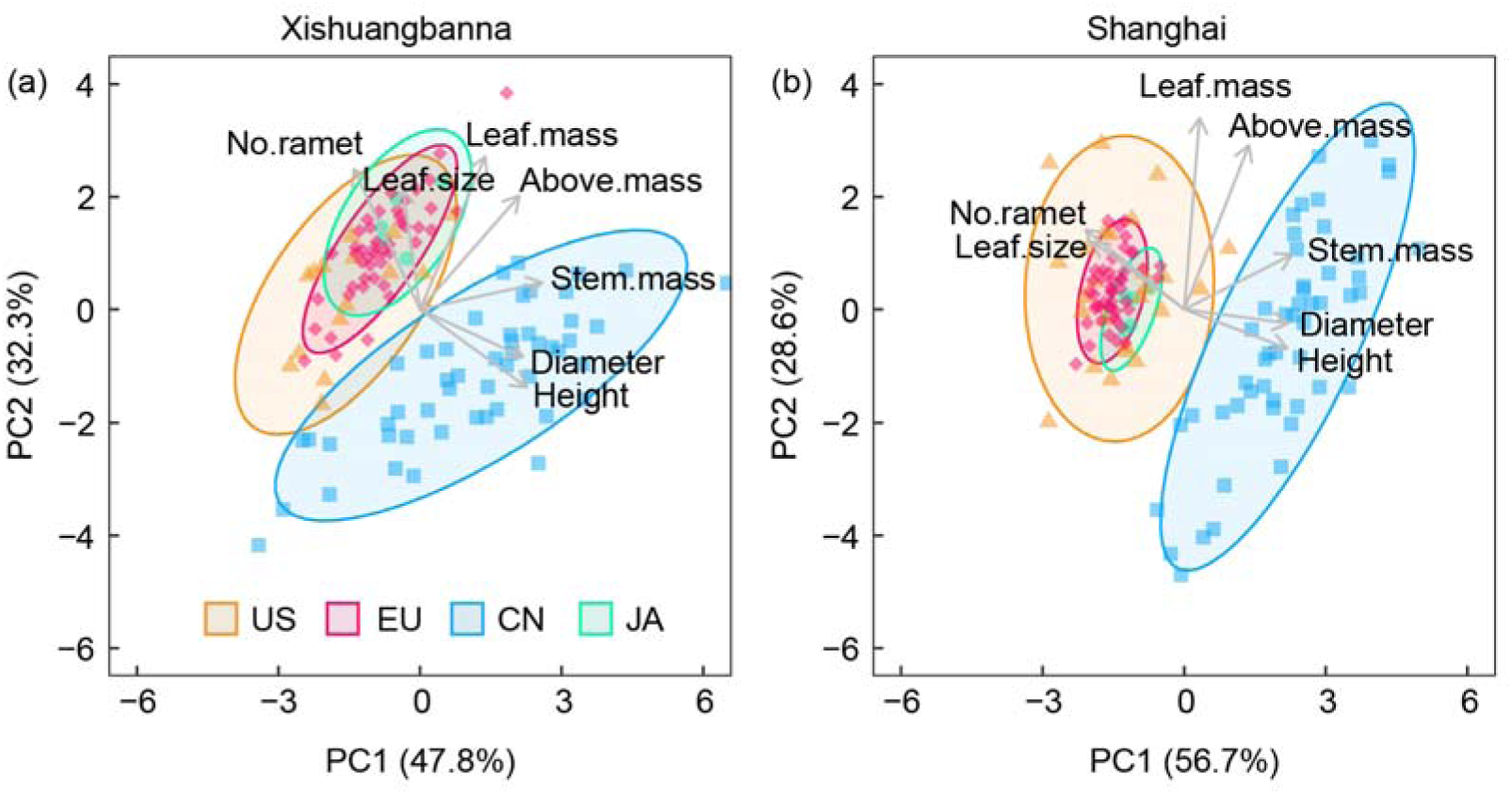
Principal components analysis of plant traits of *R. japonica* measured in the Xishuangbanna (a) and Shanghai (b) common gardens. Plant trait loading vectors are plotted as arrows. Stem.mass: stem biomass; Leaf.mass: leaf biomass; Above.mass: aboveground biomass; US: introduced North America; EU: introduced Europe; CN: native China; JA: native Japan.

The separation of Chinese populations from those from Japan, North America and Europe was recapitulated in analyses of trait responses. Plants from the native Japanese populations were not significantly different from those from introduced North American and European populations in mean for most traits (except basal diameter in Shanghai; Figure 3d). Japanese and introduced plants were shorter and thinner but had more ramets than Chinese plants (Figure 3a-f). We found no significant differences in aboveground biomass among ranges (Figure 3g, h).

**Figure 3.**
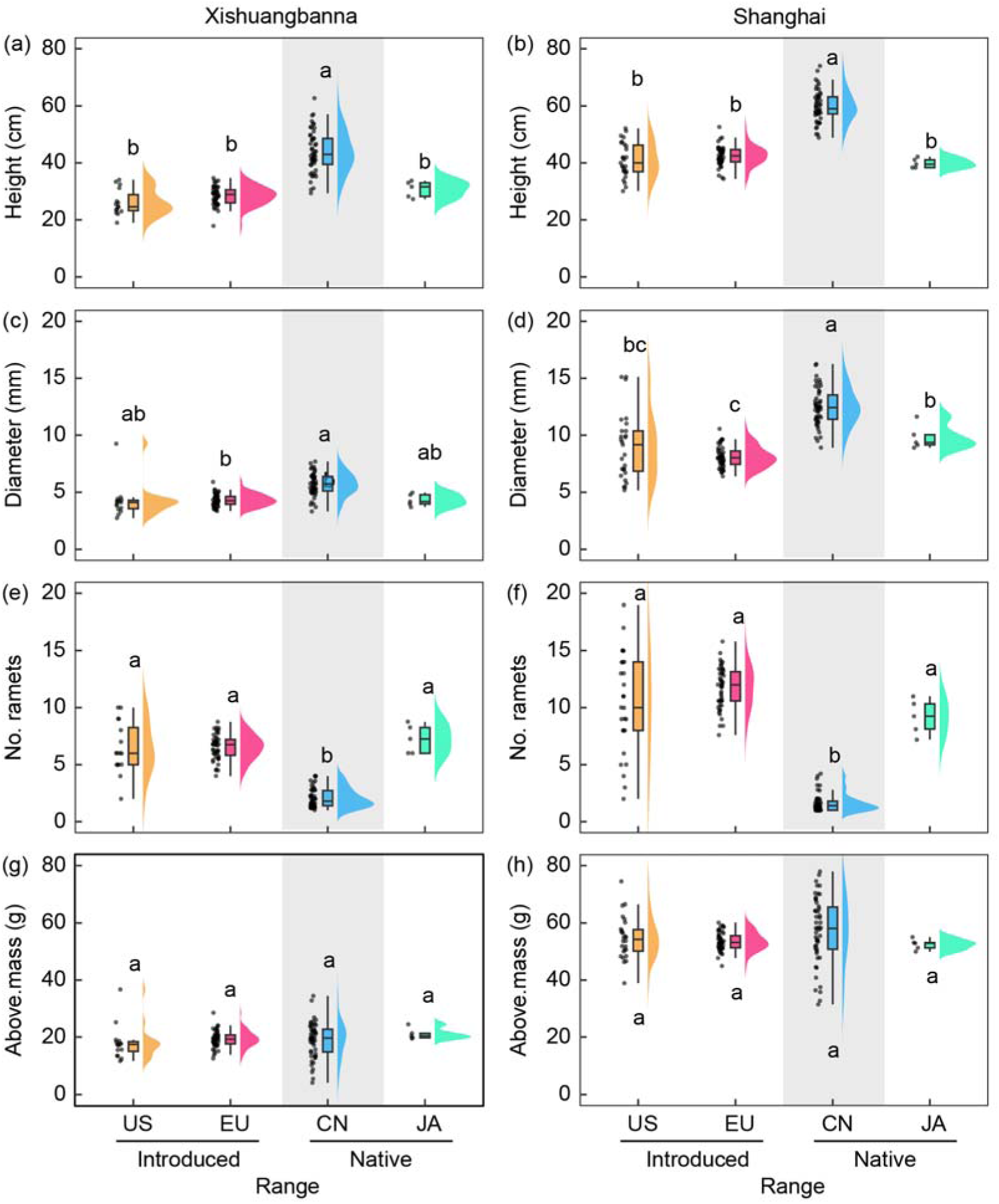
Variation in plant traits of *R. japonica* from introduced North American (US) and European (EU) and native Chinese (CN) and Japanese (JA) ranges in Xishuangbanna (a, c, e, g) and Shanghai (b, d, f, h) common gardens. Graphs are represented by population mean values (points), box with hinges and whiskers, and violin plot based on Kernel density function. Different letters represent significant differences in plant trait values among ranges. Above.mass: aboveground biomass.

### 3.2 Divergence in phenotypic plasticity

We found significant range by garden effects for almost all traits (Table 1, Table S2). Plasticity in plant height, basal diameter and aboveground biomass was lower for the Japanese and introduced populations compared to Chinese populations (Figure 4a, b, d). Japanese and introduced plants had higher plasticity in number of ramets compared to the native Chinese populations, and introduced populations had higher plasticity than those from Japan (Figure 4c).

**Figure 4.**
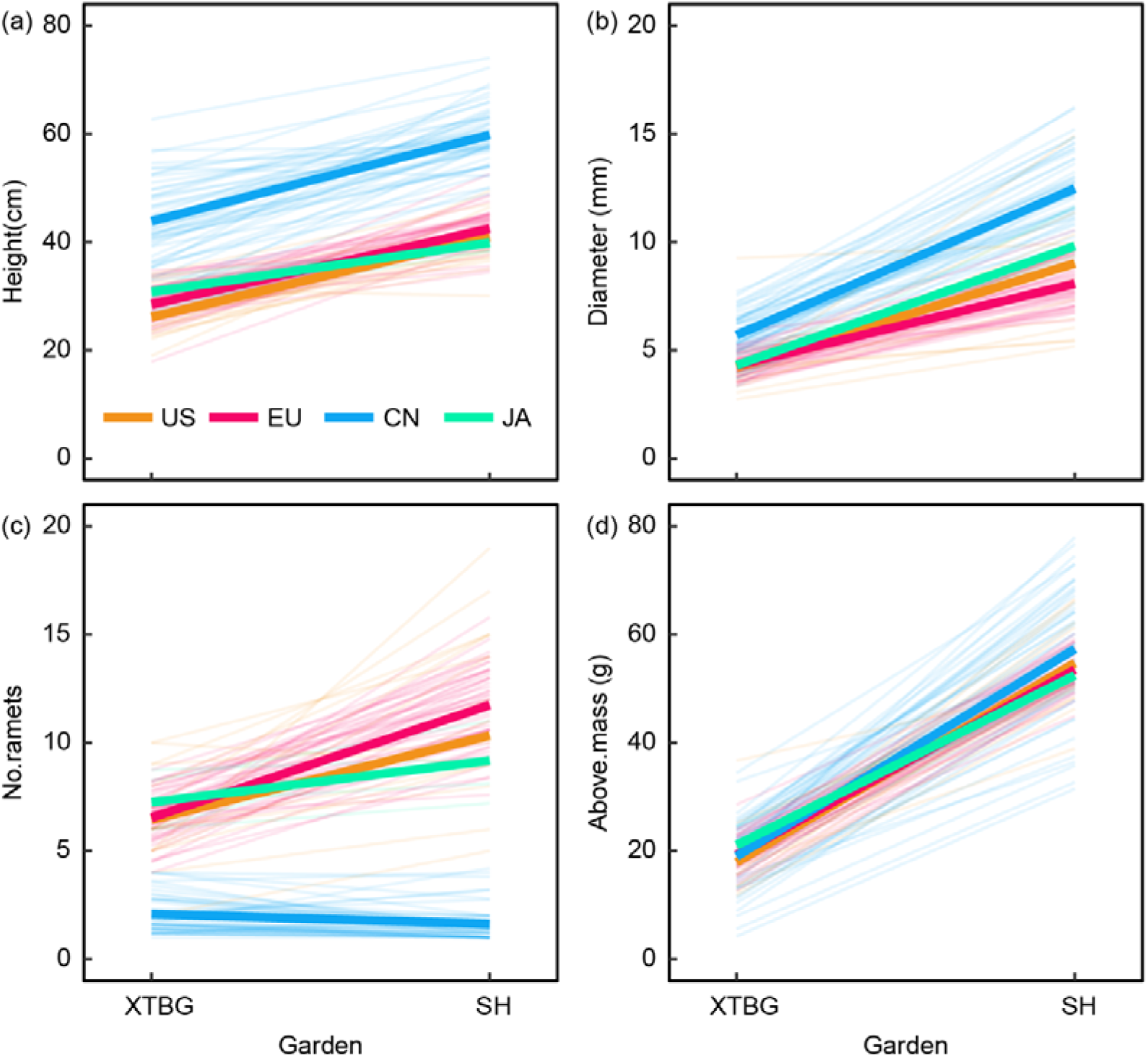
Reaction norms for populations of *R. japonica* from introduced North American (US) and European (EU) and native Chinese (CN) and Japanese (JA) ranges grown under Xishuangbanna (XTBG) and Shanghai (SH) common gardens. Mean response for each population is indicated by thin lines. Mean response for each range is indicated by thick lines. Above.mass: aboveground biomass. See more details of statistical significance in Table 1.

## 4 DISCUSSION

Introduced species around the globe provide unique opportunities to examine mechanisms that allow for rapid response to novel and changing environmental conditions (Lee, 2002; Estoup et al., 2016). Many different mechanisms—including introduction of “pre-adapted” genotypes, and the evolution of plant traits in response to novel environments—have been examined to explain the biogeographic differences of invasive species (Callaway et al., 2022; Hierro et al., 2022; Gioria et al. 2023). In previous work, we found that in the field plants in introduced *Reynoutria japonica* populations were larger with higher nutritional value compared to conspecific in native populations (Irimia et al., 2024). Our results here showed that when grown in common gardens, the same *R. japonica* plants we sampled in the introduced range have not evolved larger sizes, which does not support our first hypothesis. In fact, we found that plants from all four ranges had equivalent aboveground biomass in both gardens. However, we found support for our second hypothesis because introduced plants were not differentiated from those from the putative source native Japanese populations in most traits. Plants from China were taller and had fewer ramets than plants from Japan, North America and Europe, reflecting broad differences in the native range of this species. We also found support for our third hypothesis related to evolution of plasticity because we were able to evaluate plants from the source native populations of Japan separately from the native populations in China. Both introduced North American and European populations were more plastic in number of ramets than putative source populations in response to our common garden sites, indicating a post-introduction evolution of higher clonality in the introduced ranges.

Our results suggested that the introduction of a highly clonal genotype (general purpose genotype) from Japan might have contributed to the invasion success of *R. japonica*. In our experiment, we found that introduced plants consistently produced many more ramets than Chinese populations but were similar to Japanese populations. This finding is in line with reports that the strong vegetative reproductive ability of introduced *R. japonica* underlies its expansion into more habitats in its introduced ranges (Beerling et al., 1994; de Waal, 2001). In general, native Chinese populations were taller and produced fewer ramets than introduced and native Japanese genotypes, which might reflect an evolutionary history that favors height growth over clonal reproduction in the Chinese populations (Bossdorf et al., 2004; van Kleunen et al., 2001).

Phenotypic plasticity is thought to be useful to founding populations by extending niche breadth (Baker, 1965; Bossdorf et al., 2005; Richards et al., 2006), and could be important for adaptation to global change and increased unpredictability of climatic anomalies (Nicotra et al., 2010; Vázquez et al., 2017). However, phenotypic plasticity is the property of specific traits and its importance can vary across traits and environmental contexts (Richards et al., 2006). We found large phenotypic changes in response to our common garden sites, but Chinese plants were more plastic in plant height, basal diameter and aboveground biomass than introduced or Japanese plants. We also found higher plasticity in basal diameter in plants from Japan than those from the introduced range in North America and Europe. The results suggest that plasticity in these traits may not provide an advantage in the introduced range.

Nevertheless, our experiment did support the plasticity-first hypothesis. We showed that the introduced plants exhibited traits that were not different from those of the presumed source of the introduction (Japanese populations) in each of the two common gardens. The introduced plants were also not different from Japanese plants in plasticity of most traits. However, our results supported increased plasticity in number of ramets from both introduced ranges compared to the native Japanese populations. In sum, our findings provided evidence that an evolutionary shift to increased plasticity in clonality may have facilitated the invasion success of *R. japonica* (Levis & Pfennig, 2016).

Considering the size and ploidy levels of the knotweed genome, plasticity to novel conditions in the introduced range may have exposed cryptic genetic variation that was not expressed in the native range (i.e., the so-called “plasticity first” hypothesis; Levis & Pfennig, 2016; Mounger et al., 2021). Then, selection could act on this variation and increase standing levels of plasticity in the introduced populations. Another study found support for increased ability to respond specifically to well-watered conditions by producing more clonal propagules in *Helianthus tuberosus* (Bock et al., 2018). They found evidence for hybrid vigor in the introduced populations and two specific quantitative trait loci (QTL) that were associated with the increased ability to respond to water content in the introduced habitat. On the other hand, several studies have reported that somatic mutation may allow asexual species to maintain abundant genetic variation and adapt to changing environmental conditions (reviewed in Schoen & Schultz, 2019; see also discussions in Chen et al., 2020; Robertson et al., 2020). Further work in *R. japonica* is required to link specific genomic changes in this species that allowed for this increase in clonality.

We acknowledge that these findings are preliminary given some of the limitations in our study. For example, plants from the four ranges accumulated similar aboveground biomass, implying two distinct trait strategies to achieve the same level of fitness in the common garden environments during this growing season. Many individuals may not flower at all in the field, but persist and spread from year to year so biomass is an important indicator of fitness (Yuan et al., 2024). However, our study did not directly relate these traits to competitive ability (e. g., competitive suppression or competitive tolerance). In addition, the common garden sites in our experimental set-up could represent extreme conditions for some populations, especially in the XTBG experimental sites in southern China. Ideally, we would compare common gardens of the full collection of native and introduced *R. japonica* in North America and Europe to evaluate the conditions in the introduced range, but, due to ethical constraints, it is not feasible to establish such gardens. Finally, although we have some evidence of history and phylogenetic relationships of the plants we used, we collected a limited number of Japanese populations and could not explicitly verify the source population in Japan. Despite these limitations, we elicited plant trait differences among four ranges across the two common gardens, which provides insight into the biology of this globally invasive species. Further work is needed to confirm whether the differences we identified have contributed to the competitive advantage of the introduced populations (Alexander & Levine, 2019).

## 5 CONCLUSIONS

Although comparisons of plant traits between native and introduced populations have been widely used to examine the rapid evolution of invasive plants, the differences in growth and reproduction traits between introduced plants and those from source and non-source native populations have seldom been tested at the same time. Our study allowed us to examine evolutionary changes in trait means and plasticities in two distinct common garden environments. A novel finding is that the trait expressions in growth and reproduction of introduced plants were very similar to those from the putative source populations, but not other native populations. Introduced populations may not express higher growth performance than all native populations, which would challenge the idea that evolution of increased competitive ability is universal in invasive plants. In the case of *R. japonica*, plants from the source native and introduced populations also exhibit higher plasticity in clonality than non-source native populations, suggesting that the introduction of general-purpose genotypes may have enhanced the potential for establishment and invasion into wider ranges. Moreover, our results show that the evolutionary shift of higher plasticity in clonality, but not other traits, may further facilitate the large-scale colonization of *R. japonica* in introduced ranges. Our ability to identify the potential role of clonality and plasticity of these plants emphasizes the importance of combining the invasion history of widely distributed native and introduced populations when evaluating evolutionary responses (Hierro et al., 2005; Colautti & Lau, 2015).

## Supporting information

This

## Acknowledgements

We thank Fatima Elkott, Christiane Karasch-Wittmann, Elodie Kugler, Rongjin Li, Jeannie Mounger, Julia Rafalski, Conner Richardson, Eva Schloter, Sabine Silberhorn, Weihan Zhao, Wenchao Zhong, Zhiming Zhong and Xin Zhuang for help with field collections, and Wenbian Bo, Jieren Jin, Guangye Li, Shibo Tian, Chaonan Wang, Jiawei Wang, lei Wang and Niyao Xiang with common garden experiments. We also thank Center for Gardening and Horticulture, Seed Bank of tropical plant resources, Xishuangbanna Tropical Botanical Garden, Chinese Academy of Sciences for providing quarantine facilities and cold rooms. This study was supported by the National Natural Science Foundation of China (31961133028), the German Federal Ministry of Education and Research (MOPGA Project 306055), the German Research Foundation (grant 431595342) and the Chinese Academy of Sciences President’s International Fellowship Initiative (2023VBB0012).

## Competing interests

The authors declare no conflict of interest.

## Data Availability Statement

The data that support the findings of this study are openly available in Dryad at http://doi.org/10.6084/m9.figshare.27216120.

## Author contributions

**Shengyu Wang:** Conceptualization; data curation; formal analysis; investigation; methodology; project administration; resources; validation; visualization; writing – original draft preparation; writing – review and editing. **Zhiyong Liao:** Conceptualization; data curation; formal analysis; investigation; methodology; project administration; resources; validation; visualization; writing – review and editing. **Peipei Cao**: Investigation; project administration; resources; writing – review and editing; **Marc W. Schmid**: Formal analysis; methodology; writing – review and editing; **Lei Zhang**: Investigation; resources; writing – review and editing. **Jingwen Bi**: Investigation; resources; writing – review and editing. **Stacy B. Endriss:** Writing – review and editing. **Yujie Zhao:** Investigation; resources; writing – review and editing. **Madalin Parepa:** Visualization; writing – review and editing. **Wenyi Hu:** Resources, writing – review and editing. **Hikaru Akamine:** Resources, writing – review and editing. **Jihua Wu:** Conceptualization; review and editing. **Ruiting Ju:** Conceptualization; review and editing. **Oliver Bossdorf:** Conceptualization; funding acquisition; project administration; review and editing. **Christina L. Richards:** Conceptualization; funding acquisition; project administration; review and editing; supervision. **Bo Li:** Conceptualization; funding acquisition; project administration; review and editing; supervision.

## Notes

### Competing Interest Statement

The authors have declared no competing interest.

