## Supplementary material for "General purpose genotypes and evolution of higher plasticity in clonality underlie knotweed invasion": This

**Supplemental materials**

**Table	S1** Geographic origins of the 128 studied populations of *Reynoutria japonica* and their uses in the Xishuangbanna Tropical Botanical Garden (XTBG) and Shanghai (SH) common gardens.

| Range | Population ID | Garden | Latitude | Longitude |
| --- | --- | --- | --- | --- |
| US | US01 | XTBG/SH | 34.24 | -83.46 |
| US | US04 | XTBG/SH | 35.1 | -83.1 |
| US | US06 | XTBG/SH | 35.74 | -82.68 |
| US | US08 | XTBG/SH | 36.21 | -81.78 |
| US | US09 | XTBG/SH | 36.38 | -81.38 |
| US | US10 | XTBG/SH | 36.65 | -80.92 |
| US | US11 | SH | 36.67 | -80.57 |
| US | US14 | SH | 37.37 | -79.41 |
| US | US15 | SH | 37.43 | -79.16 |
| US | US18 | XTBG/SH | 38.46 | -78.59 |
| US | US19 | SH | 38.65 | -78.54 |
| US | US20 | XTBG/SH | 39.08 | -78.09 |
| US | US22 | XTBG/SH | 39.61 | -76.68 |
| US | US31 | XTBG/SH | 41.01 | -74.35 |
| US | US33 | XTBG/SH | 41.41 | -73.96 |
| US | US34 | SH | 41.77 | -73.86 |
| US | US38 | XTBG/SH | 42.64 | -72.91 |
| US | US39 | XTBG/SH | 42.92 | -72.77 |
| US | US40 | XTBG/SH | 43.2 | -72.45 |
| US | US42 | SH | 43.84 | -72.19 |
| US | US43 | XTBG/SH | 43.94 | -71.68 |
| US | US44 | XTBG/SH | 44.12 | -71.18 |
| US | US45 | SH | 44.11 | -70.69 |
| US | US46 | SH | 44.21 | -70.31 |
| US | US47 | SH | 44.53 | -69.89 |
| US | US49 | SH | 44.94 | -68.99 |
| US | US50 | SH | 44.95 | -68.64 |
| EU | EU01 | XTBG/SH | 44.88 | 7.69 |
| EU | EU02 | XTBG/SH | 44.75 | 7.48 |
| EU | EU03 | XTBG/SH | 44.67 | 7.29 |
| EU | EU04 | XTBG/SH | 45.19 | 8.04 |
| EU | EU06 | XTBG/SH | 45.64 | 8.37 |
| EU | EU07 | XTBG/SH | 45.8 | 8.87 |
| EU | EU09 | XTBG/SH | 46.27 | 9 |
| EU | EU10 | XTBG/SH | 46.51 | 8.7 |
| EU | EU11 | XTBG/SH | 47.16 | 8.56 |
| EU | EU12 | XTBG/SH | 47.62 | 8.22 |
| EU | EU13 | XTBG/SH | 47.81 | 7.61 |
| EU | EU14 | XTBG/SH | 47.94 | 7.88 |
| EU | EU15 | XTBG/SH | 48.28 | 8.11 |
| EU | EU16 | XTBG/SH | 48.37 | 8.56 |
| EU | EU17 | XTBG/SH | 48.47 | 8.92 |
| EU | EU18 | XTBG/SH | 48.56 | 8.39 |
| EU | EU19 | XTBG/SH | 48.79 | 8.32 |
| EU | EU21 | XTBG/SH | 49.59 | 8.73 |
| EU | EU22 | XTBG/SH | 49.9 | 8.83 |
| EU | EU23 | XTBG/SH | 50.07 | 8.48 |
| EU | EU25 | XTBG/SH | 50.31 | 7.79 |
| EU | EU26 | XTBG/SH | 50.54 | 7.08 |
| EU | EU27 | XTBG/SH | 51.14 | 6.78 |
| EU | EU28 | XTBG/SH | 51.43 | 7.29 |
| EU | EU29 | XTBG/SH | 51.87 | 7.55 |
| EU | EU30 | XTBG/SH | 52.15 | 7.62 |
| EU | EU31 | XTBG/SH | 52.39 | 7.94 |
| EU | EU32 | XTBG/SH | 52.72 | 8.26 |
| EU | EU33 | XTBG/SH | 53.01 | 8.7 |
| EU | EU34 | XTBG/SH | 53.22 | 9.57 |
| EU | EU35 | XTBG/SH | 53.45 | 10.08 |
| EU | EU36 | XTBG/SH | 53.99 | 10.25 |
| EU | EU37 | XTBG/SH | 54.31 | 10.13 |
| EU | EU38 | XTBG/SH | 54.71 | 11.45 |
| EU | EU39 | XTBG/SH | 55.46 | 12.19 |
| EU | EU40 | XTBG/SH | 55.68 | 13.19 |
| EU | EU41 | XTBG/SH | 56.14 | 13.76 |
| EU | EU42 | XTBG/SH | 56.45 | 13.6 |
| EU | EU43 | XTBG/SH | 56.83 | 13.96 |
| EU | EU44 | XTBG/SH | 57.7 | 14.11 |
| EU | EU45 | XTBG/SH | 58.17 | 14.58 |
| EU | EU46 | XTBG/SH | 58.41 | 15.65 |
| EU | EU47 | XTBG/SH | 58.67 | 16.2 |
| EU | EU48 | XTBG/SH | 58.89 | 17.56 |
| EU | EU49 | XTBG/SH | 59.32 | 18.02 |
| EU | EU50 | XTBG/SH | 59.95 | 17.71 |
| CN | CN01 | XTBG/SH | 29.31 | 117.85 |
| CN | CN02 | XTBG/SH | 29.21 | 117.15 |
| CN | CN03 | XTBG/SH | 28.71 | 117.02 |
| CN | CN04 | XTBG/SH | 28.54 | 117.05 |
| CN | CN05 | XTBG/SH | 28.19 | 116.76 |
| CN | CN06 | XTBG/SH | 27.87 | 116.78 |
| CN | CN07 | XTBG/SH | 27.66 | 116.68 |
| CN | CN08 | XTBG/SH | 27.38 | 116.45 |
| CN | CN09 | XTBG/SH | 27.08 | 116.34 |
| CN | CN10 | XTBG/SH | 26.85 | 116.37 |
| CN | CN11 | XTBG/SH | 26.55 | 116.27 |
| CN | CN12 | XTBG/SH | 26.27 | 116.32 |
| CN | CN13 | XTBG/SH | 26.03 | 116.16 |
| CN | CN14 | XTBG/SH | 25.8 | 115.94 |
| CN | CN15 | XTBG/SH | 25.53 | 115.84 |
| CN | CN16 | XTBG/SH | 25.24 | 115.73 |
| CN | CN17 | XTBG/SH | 24.96 | 115.61 |
| CN | CN18 | XTBG/SH | 24.71 | 115.82 |
| CN | CN19 | XTBG/SH | 23.82 | 115.38 |
| CN | CN20 | XTBG/SH | 23.75 | 115.27 |
| CN | CN21 | XTBG/SH | 23.64 | 113.84 |
| CN | CN22 | XTBG/SH | 23.29 | 114.01 |
| CN | CN23 | XTBG/SH | 23.74 | 113.91 |
| CN | CN24 | XTBG/SH | 24.44 | 113.25 |
| CN | CN25 | XTBG/SH | 24.62 | 113.76 |
| CN | CN26 | XTBG/SH | 26.1 | 114.69 |
| CN | CN27 | XTBG/SH | 29.1 | 117.12 |
| CN | CN28 | XTBG/SH | 29.43 | 117.16 |
| CN | CN29 | XTBG/SH | 29.64 | 117.22 |
| CN | CN30 | XTBG/SH | 29.81 | 117.05 |
| CN | CN31 | XTBG/SH | 29.98 | 116.97 |
| CN | CN32 | XTBG/SH | 30.18 | 117.04 |
| CN | CN33 | XTBG/SH | 30.44 | 116.23 |
| CN | CN34 | XTBG/SH | 30.69 | 115.98 |
| CN | CN35 | XTBG/SH | 30.82 | 116.05 |
| CN | CN36 | XTBG/SH | 30.92 | 116.21 |
| CN | CN37 | XTBG/SH | 31.09 | 116.1 |
| CN | CN38 | XTBG/SH | 31.21 | 116.02 |
| CN | CN39 | XTBG/SH | 31.4 | 115.93 |
| CN | CN40 | XTBG/SH | 31.58 | 115.97 |
| CN | CN41 | XTBG/SH | 31.81 | 115.85 |
| CN | CN42 | XTBG/SH | 34.27 | 115.68 |
| CN | CN43 | XTBG/SH | 34.72 | 116.79 |
| CN | CN44 | XTBG/SH | 35.06 | 116.31 |
| CN | CN45 | XTBG/SH | 35.52 | 117.82 |
| CN | CN46 | XTBG/SH | 35.74 | 117.64 |
| CN | CN47 | XTBG/SH | 36 | 117.65 |
| CN | CN48 | XTBG/SH | 36.38 | 117.66 |
| CN | CN49 | XTBG/SH | 36.87 | 117.68 |
| CN | CN50 | XTBG/SH | 32.85 | 118.44 |
| JA | JA02 | XTBG/SH | 32.81 | 129.92 |
| JA | JA03 | XTBG/SH | 32.99 | 129.77 |
| JA | JA04 | XTBG/SH | 33.21 | 129.68 |
| JA | JA05 | XTBG/SH | 33.32 | 129.6 |
| JA | JA06 | XTBG/SH | 33.33 | 129.53 |

**Table S2** The effects of range, garden and their interactions on plant traits of *R. japonica*. Df, Degrees of freedom; SS, Sum of squares. P-values were calculated by treating population as a random factor. Thus, the factor range and its contrasts were tested against population, and the interactions of range and its contrasts with garden against garden:population. Garden was tested against the residuals. Even though population was treated as a random factor, the variation explained by it and the results of a test against residuals were provided.

|  |  | Height | | | |  | Diameter | | | |
| --- | --- | --- | --- | --- | --- | --- | --- | --- | --- | --- |
| Term |  | Df | SS | P | SS% |  | Df | SS | P | SS% |
| Garden |  | 1 | 46890.53 | 0.000 | 26.73 |  | 1 | 6762.6 | 0.000 | 53.69 |
| Range, contrasts: |  | 3 | 69273.57 | 0.000 | 39.49 |  | 3 | 1996.59 | 0.000 | 15.85 |
| a) CN vs. JA/EU/US |  | 1 | 69083.93 | 0.000 | 39.38 |  | 1 | 1939.93 | 0.000 | 15.4 |
| b) JA vs. EU/US |  | 1 | 43.12 | 0.571 | 0.02 |  | 1 | 49.98 | 0.014 | 0.4 |
| c) EU vs. US |  | 1 | 146.52 | 0.297 | 0.08 |  | 1 | 6.69 | 0.363 | 0.05 |
| Population |  | 124 | 16536.91 | 0.000 | 9.43 |  | 124 | 993.39 | 0.000 | 7.89 |
| Garden:Range, contrasts: |  | 3 | 489.33 | 0.101 | 0.28 |  | 3 | 485.35 | 0.000 | 3.85 |
| a) Garden:CN vs. JA/EU/US |  | 1 | 329.18 | 0.041 | 0.19 |  | 1 | 446.86 | 0.000 | 3.55 |
| b) Garden:JA vs. EU/US |  | 1 | 160.11 | 0.151 | 0.09 |  | 1 | 36.79 | 0.001 | 0.29 |
| c) Garden:EU vs. US |  | 1 | 0.038 | 0.982 | 0 |  | 1 | 1.71 | 0.473 | 0.01 |
| Garden:Population |  | 112 | 8584.43 | 0.000 | 4.89 |  | 112 | 368.87 | 0.071 | 2.93 |
| Residuals |  | 738 | 33664.31 |  | 19.19 |  | 738 | 1987.84 |  | 15.78 |
|  |  | Leaf size | | | |  | Leaf biomass | | | |
| Term |  | Df | SS | P | SS% |  | Df | SS | P | SS% |
| Garden |  | 1 | 113061.3 | 0.000 | 4.18 |  | 1 | 110185.8 | 0.000 | 78 |
| Range, contrasts: |  | 3 | 842499.8 | 0.000 | 31.15 |  | 3 | 716.36 | 0.080 | 0.51 |
| a) CN vs. JA/EU/US |  | 1 | 830797.1 | 0.000 | 30.72 |  | 1 | 712.51 | 0.010 | 0.5 |
| b) JA vs. EU/US |  | 1 | 97.58 | 0.886 | 0 |  | 1 | 0.0527 | 0.982 | 0 |
| c) EU vs. US |  | 1 | 11605.2 | 0.119 | 0.43 |  | 1 | 3.8 | 0.848 | 0 |
| Population |  | 124 | 583834.3 | 0.000 | 21.59 |  | 123 | 12729.1 | 0.000 | 9.01 |
| Garden:Range, contrasts: |  | 3 | 57697.15 | 0.000 | 2.13 |  | 3 | 107.92 | 0.402 | 0.08 |
| a) Garden:CN vs. JA/EU/US |  | 1 | 47256.44 | 0.000 | 1.75 |  | 1 | 40.84 | 0.292 | 0.03 |
| b) Garden:JA vs. EU/US |  | 1 | 654.14 | 0.616 | 0.02 |  | 1 | 41.13 | 0.291 | 0.03 |
| c) Garden:EU vs. US |  | 1 | 9786.56 | 0.054 | 0.36 |  | 1 | 25.94 | 0.401 | 0.02 |
| Garden:Population |  | 113 | 292404.7 | 0.000 | 10.81 |  | 113 | 4124.01 | 0.000 | 2.92 |
| Residuals |  | 735 | 815056.3 |  | 30.14 |  | 707 | 13404.27 |  | 9.49 |
|  |  | Stem biomass | | | |  | Aboveground biomass | | | |
| Term |  | Df | SS | P | SS% |  | Df | SS | P | SS% |
| Garden |  | 1 | 31934.59 | 0.000 | 64.31 |  | 1 | 309293.3 | 0.000 | 81.17 |
| Range, contrasts: |  | 3 | 4559.08 | 0.000 | 9.18 |  | 3 | 930.14 | 0.340 | 0.24 |
| a) CN vs. JA/EU/US |  | 1 | 4549.75 | 0.000 | 9.16 |  | 1 | 929.04 | 0.068 | 0.24 |
| b) JA vs. EU/US |  | 1 | 2.49 | 0.815 | 0.01 |  | 1 | 1.07 | 0.950 | 0 |
| c) EU vs. US |  | 1 | 6.83 | 0.699 | 0.01 |  | 1 | 0.0346 | 0.991 | 0 |
| Population |  | 123 | 5589.49 | 0.000 | 11.26 |  | 123 | 33737.48 | 0.000 | 8.85 |
| Garden:Range, contrasts: |  | 3 | 1970.66 | 0.000 | 3.97 |  | 3 | 1301.2 | 0.002 | 0.34 |
| a) Garden:CN vs. JA/EU/US |  | 1 | 1970.48 | 0.000 | 3.97 |  | 1 | 1198.04 | 0.000 | 0.31 |
| b) Garden:JA vs. EU/US |  | 1 | 0.11 | 0.928 | 0 |  | 1 | 86.36 | 0.311 | 0.02 |
| c) Garden:EU vs. US |  | 1 | 0.0713 | 0.943 | 0 |  | 1 | 16.79 | 0.654 | 0 |
| Garden:Population |  | 113 | 1556.04 | 0.000 | 3.13 |  | 113 | 9403.32 | 0.000 | 2.47 |
| Residuals |  | 716 | 4050.59 |  | 8.16 |  | 707 | 26379.79 |  | 6.92 |
|  |  | No. ramets | | | |  |  |  |  |  |
| Term |  | Df | SS | P | SS% |  |  |  |  |  |
| Garden |  | 1 | 1504.7 | 0.000 | 5.97 |  |  |  |  |  |
| Range, contrasts: |  | 3 | 12939.17 | 0.000 | 51.33 |  |  |  |  |  |
| a) CN vs. JA/EU/US |  | 1 | 12888.86 | 0.000 | 51.13 |  |  |  |  |  |
| b) JA vs. EU/US |  | 1 | 33.66 | 0.073 | 0.13 |  |  |  |  |  |
| c) EU vs. US |  | 1 | 16.66 | 0.206 | 0.07 |  |  |  |  |  |
| Population |  | 124 | 1278.84 | 0.302 | 5.07 |  |  |  |  |  |
| Garden:Range, contrasts: |  | 3 | 1849.54 | 0.000 | 7.34 |  |  |  |  |  |
| a) Garden:CN vs. JA/EU/US |  | 1 | 1677.11 | 0.000 | 6.65 |  |  |  |  |  |
| b) Garden:JA vs. EU/US |  | 1 | 172.41 | 0.000 | 0.68 |  |  |  |  |  |
| c) Garden:EU vs. US |  | 1 | 0.0197 | 0.949 | 0 |  |  |  |  |  |
| Garden:Population |  | 113 | 532.29 | 1.000 | 2.11 |  |  |  |  |  |
| Residuals |  | 736 | 7104.38 |  | 28.18 |  |  |  |  |  |

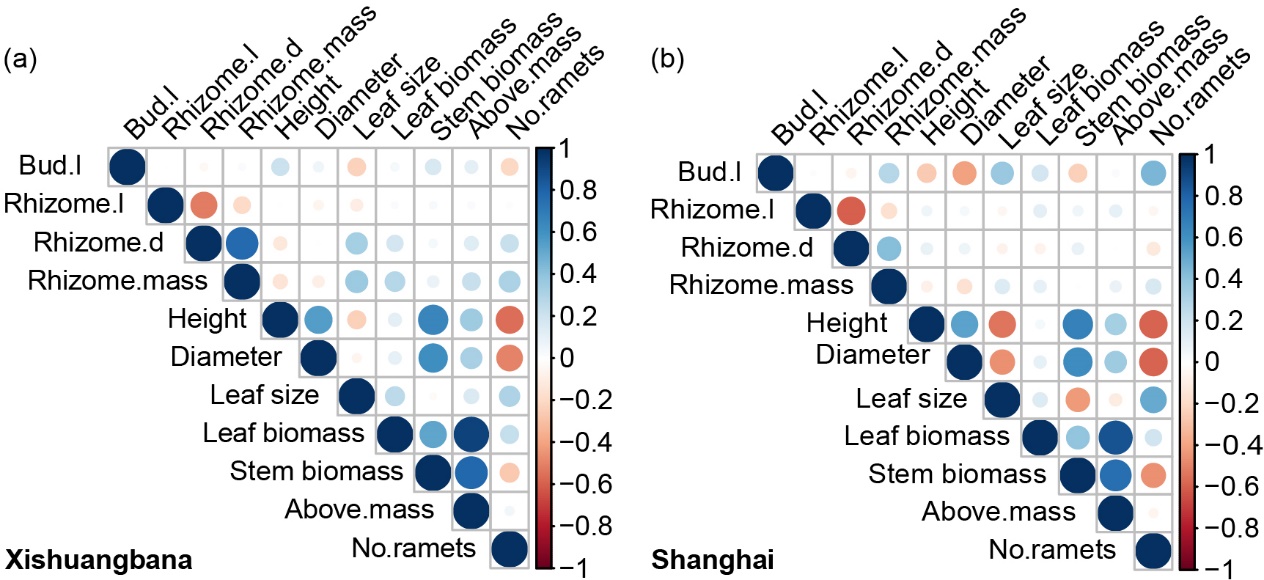

**Figure S1** Pearson correlation coefficient for initial rhizome size and traits of *R. japonica* grown in the Xishuangbanna and Shanghai common gardens. Bul.l: length of bud; Rhizome.l: length of rhizome; Rhizome.d: diameter of rhizome; Rhizome.mass: fresh mass of rhizome; Above.mass: aboveground biomass.
